# Divergent evolution of genetic sex determination mechanisms along environmental gradients

**DOI:** 10.1101/2022.05.23.493175

**Authors:** Martijn A. Schenkel, Jean-Christophe Billeter, Leo W. Beukeboom, Ido Pen

## Abstract

Sex determination (SD) is a crucial developmental process, but its molecular underpinnings are very diverse, both between and within species. SD mechanisms have traditionally been categorized as either genetic (GSD) or environmental (ESD), depending on the type of cue that triggers sexual differentiation. However, mixed systems, with both genetic and environmental components, are more prevalent than previously thought. Here, we show theoretically that environmental effects on expression levels of genes within SD regulatory mechanisms can easily trigger within-species evolutionary divergence of SD mechanisms. This may lead to the stable coexistence of multiple SD mechanisms and to spatial variation in the occurrence of different SD mechanisms along environmental gradients. We applied the model to the SD system of the housefly, a global species with world-wide latitudinal clines in the frequencies of different SD systems, and found that it correctly predicted these clines if specific genes in the housefly SD system were assumed to have temperature-dependent expression levels. We conclude that environmental sensitivity of gene regulatory networks may play an important role in diversification of SD mechanisms.

## Introduction

Sex determination (SD) is a crucial aspect of the development of sexually-reproducing organisms, yet the regulatory mechanisms underlying SD are very diverse and prone to evolutionary change^1,2^. SD mechanisms have traditionally been classified as either environmental (ESD) or genetic (GSD) depending on the type of signal that initiates the determination of an individual’s sex. Under ESD, such signal include temperature, salinity, and acidity during a sensitive period in embryonic development (reviewed in ^3,4^). Under GSD, the signal is a specific gene (or set of genes) present in zygotes, leading to either male or female development, such as the male-determining *Sex-determining Region Y (SRY)* gene in mammals or *transformer* (*tra*) in many insects^5–9^. Diversification of SD mechanisms occurs via the evolution of novel SD mechanisms that replace their predecessors, a process called SD transition. SD transitions are prevalent in some taxa but not in others^1,2^, indicating variable evolvability of SD systems. What enhances the evolvability of some SD systems but not others and what causes SD transitions is still poorly understood.

One often-overlooked aspect of the evolution of SD is how environmental and genetic factors may simultaneously affect SD. Rather than being mutually exclusive, ESD and GSD could instead be considered as two extremes of a continuum^2,10,11^, with mixed systems occurring in several organismal groups, such as amphibians, fish and insects^4,10,12–15^. In such mixed systems, SD may reflect a delicate balance between genetic effects that bias the process of SD towards male or female development, counteracted by environmental effects that push the system in the opposite direction^16,17^. GSD has repeatedly evolved in species which previously had ESD (e.g., ^18^), and several theoretical models have been developed to predict when such turnovers should occur^10,19,20^. However, such models typically do not explicitly consider the underlying molecular mechanisms of sex determination. Although sex is determined by genes in species with GSD, environmental conditions can affect SD by modifying the expression levels of these genes^21,22^. Despite clear evidence that such environmental effects may perturb the action of GSD mechanisms (e.g., ^4^), their effect on the evolution of GSD systems is still unknown.

In many species, SD involves hierarchical gene regulation^23–25^, where an initial signal targets a small number of regulatory genes that in turn regulate downstream targets. Evolutionary transitions between GSD systems are thought to occur primarily by the displacement of the initial signal gene by another gene with a similar function, or by the recruitment of a new regulatory gene on top of the ancestral SD regulatory pathway^23,24,26^. Genes downstream of the top regulatory genes are considered to be more constrained in terms of evolutionary change, as mutations in such genes may interfere with their pre-existing function in regulating SD. Nonetheless, they are not fully prohibited from undergoing further evolution, and some changes may still occur^27^.

A new evolutionary framework needs to integrate the views that (1) SD is not solely environmentally or genetically determined but is to varying degrees affected by both types of cues; and (2) changes in SD cascades do not only occur at the top but may also occur via changes in the underlying genetic network. Here, we formalize this framework by developing a theoretical model of the evolution of SD systems in the presence of spatial environmental variation that affects expression of SD genes. The model is inspired by the polymorphic SD system of the housefly *Musca domestica*. In this system, temperature is likely to act as an environmental cue because (1) variation in SD systems is correlated with variation in temperature between natural populations, and (2) temperature affects SD in several *M. domestica* strains harbouring mutant SD genes (see also Box 1). Like in *M. domestica* SD, the model features two types of SD genes: an environmentally-sensitive gene *F* induces femaleness when active, and one or more *M*-genes that induce maleness by inhibiting *F* (Figure 1A, see details below). We investigate here how the (co-)evolution of *F* and *M* can yield novel SD systems under the influence of environmental sensitivity of *F*. We then use the model to explain how the multifactorial SD system of houseflies has evolved (Box 1).

**Figure 1:**
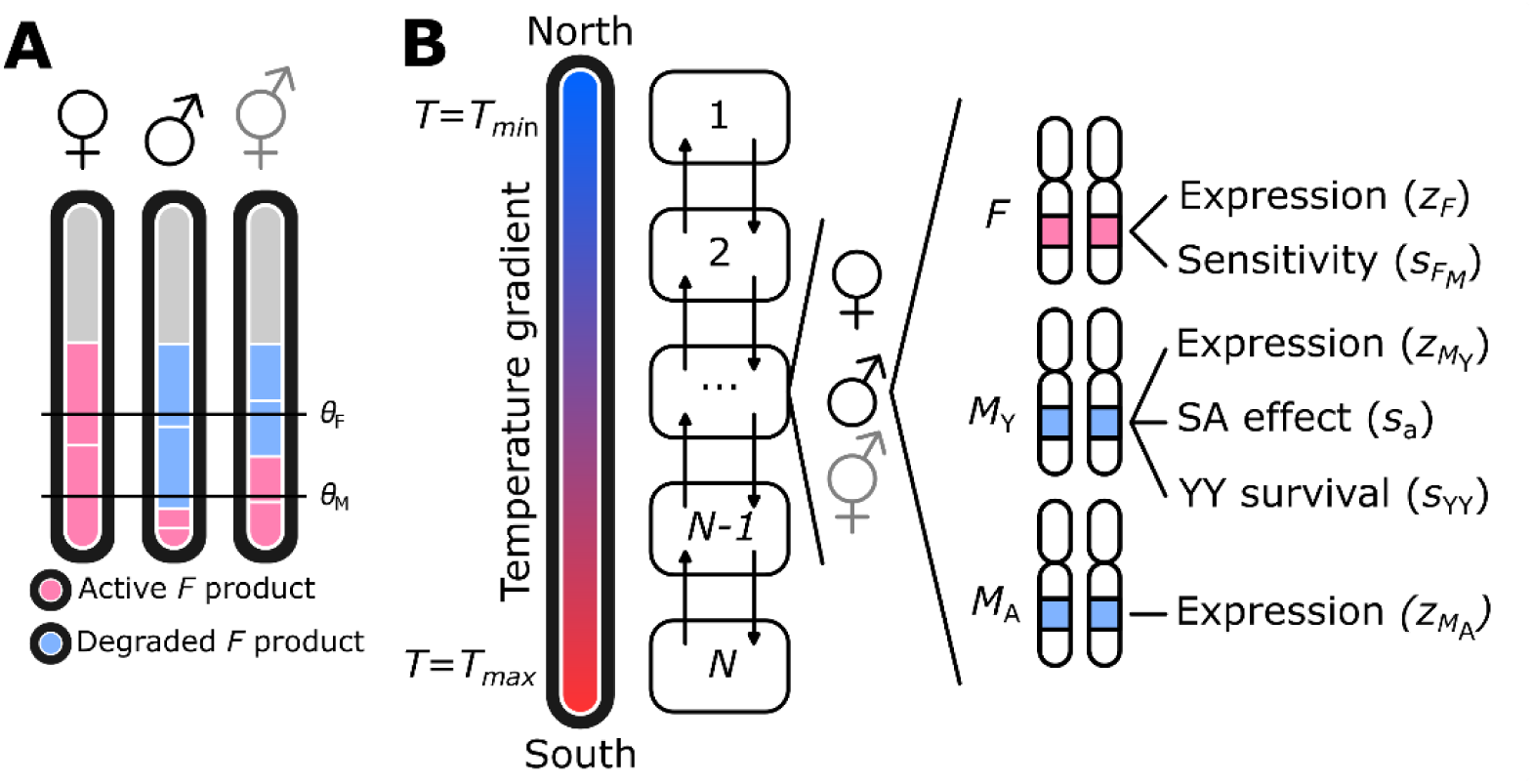
Model overview. (A) Sex is determined based on the net total active *F* product. Active *F* is produced by the *F* locus, and degraded by *M*, produced by *M*_Y_ and/or *M*_A_. Higher temperature increases expression levels of *F*. If the net total active *F* product exceeds a threshold *θ*_F_, individuals become females, whereas below the threshold *θ*_M_ individuals become males; otherwise, individuals develop into infertile intersexes. (B) Demes 1 through *N* are arranged along a linear gradient where *T* increases from *T*_min_ to *T*_max_ Each deme contains a variable number of females, males, and intersexes. Reproduction occurs by mating between males and females within the same deme; intersexes do not reproduce. Dispersal occurs at a rate *d* to neighbouring demes (indicated by arrows). Every individual has three gene loci *F, M*_Y_, and *M*_A_ that jointly determine sex.

### Model summary

Here we briefly describe the model; a more detailed description of the model and simulation techniques is in the Methods section below. We developed an individual-based simulation model, where individuals occupy a linear array of subpopulations (demes) arranged along a temperature gradient. The life cycle is as follows: adults reproduce sexually and then die; their offspring undergo sexual development and viability selection; a fraction of the surviving offspring migrates from their natal subpopulation to a neighbouring subpopulation; finally, individuals mature and the next round of reproduction begins.

Motivated by the SD mechanisms of the housefly (Box 1), individual sexual development was modelled to result from interactions between a single feminizing gene *F*, one or more masculinizing genes *M* and the local temperature of an individual’s environment. The *F* gene produces a temperature-dependent amount of product which is partially inhibited or degraded by the products from the *M* genes; the remaining or net amount of *F* product, denoted by 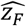, determines sex: if 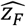 exceeds a threshold value *θ*_F_, the individual develops as female, whereas if 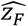 is below a second threshold value *θ*_M_ < *θ*_F_ it develops into a male. If 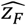 is between the two thresholds, 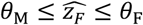, then the individual develops into an infertile intersex.

The value of 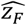 is obtained by summing up the net expression levels *z*_*F*_ of both *F* alleles in an individual. The quantitative value of *z*_*F*_ can vary between *F* alleles and depends on (1) the temperature-dependent expression level of the allele, (2) the allele’s sensitivity to *M* product, and (3) the amount of *M* product. Specifically, normalized temperature varies between *T* = 0 at one end (the “north”) of the array of subpopulations and *T* = 1 at the other end (the “south”) and it affects the net amount of *F* product according to

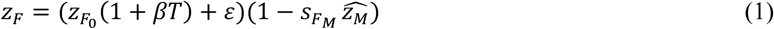

The first factor on the right, 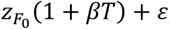, represents the initial amount of *F* product, before partial degradation by *M* product, where 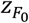is the *F* allele’s baseline expression level at *T* = 0, *β* ≥ 0 quantifies the linear rate of increase in *F* expression with temperature, and *ε* represents random variation in expression levels due to environmental and/or developmental noise. The second factor, 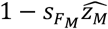, represents the proportion of *F* product that is not degraded by *M* product, where 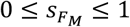 is the *F* allele’s sensitivity to *M* product and 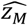 is the cumulative amount of *M* product produced by the individual’s *M* alleles.

The baseline expression level 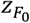and sensitivity 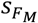 of *F* alleles are evolvable trait values, as are the expression levels of *M* alleles. Whenever a gamete is produced, each allelic trait incurs a mutation with a certain probability and its trait value is modified. Most mutations are “regular” mutations that modify traits by adding a small Gaussian amount with mean zero, but a small fraction of mutations are “null mutations” such that the resulting trait values (allelic expression levels or sensitivity to *M* product) are zero and cannot evolve any further. See Supplementary Table 2 for all model parameters and their default values used in the simulations.

The initial populations all have an XY male heterogametic system: all individuals carry two *F* alleles on an autosomal *F* locus and males additionally carry a single *M* allele (designated *M*_Y_) on their Y-chromosome. Initially there are no *M* alleles on autosomes, but we assume that during meiosis sometimes a new *M* allele (designated *M*_A_) is created on an autosomal locus; this is assumed to occur via *de novo* evolution of a novel *M* allele, but can also occur via transposition from a Y-chromosomal *M* allele. Individuals carry at most four *M* alleles: two on the Y-chromosome (if they have two Y chromosomes) and two on an autosome. In natural systems, many loci may be capable of evolving a male-determining function^28^, but we consider here only a single autosomal locus for simplicity. Thus, the initial XY population has the potential to evolve into a population with a new system of male heterogamety where an autosomal chromosome with an *M* allele has replaced the original Y-chromosome. Evolution of populations with female heterogamety is also a potential outcome, if females become heterozygous for an insensitive *F* allele (i.e., with 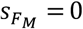) with a sufficiently high expression level.

We also allowed for Y-chromosomal fitness effects: (1) individuals homozygous for the Y-chromosome will have reduced viability 0 ≤ *s*_YY_ ≤ 1, and (2) Y-chromosomes carry sexually antagonistic alleles that are beneficial to males and detrimental in females, with additive effect 1 + *s*_*a*_ males and 1 − *s*_*a*_ in females (where *s*_*a*_ ≥ 0 is the antagonistic effect). The combined effects of Y-homozygosity and sexual antagonism are assumed to be multiplicative, i.e., a male with two Y-chromosomes will have his expected fitness modified by a factor *s*_YY_(1 + *s*_*a*_), whereas a female with two Y-chromosomes gets *s*_*YY*_ (1 − *s*_a_).

## Results

### Evolution of F insensitivity and establishment of SD system gradients

As an initial exploration of the model, we performed a set of simulations without temperature effects, where we instead varied SD thresholds *θ*_F_ and *θ*_M_ to determine how this affects the evolution of *F, M*, and the SD system as a whole. The results of this analysis are presented in detail in Appendix 1. Most importantly, we find that *F* can evolve into a dominant female-determining gene by becoming insensitive to *M*, provided that a single *F* allele generates enough *F* product to induce feminization, i.e., the threshold for feminization *θ*_F_ is sufficiently low relative to *F* expression *z*_*F*_. We refer to such insensitive dominant feminizing *F* alleles as *F*_I_. Provided that the feminization threshold *θ*_F_ is constant under all conditions, local variation in temperature could then lead to divergent evolution of SD systems as temperature effects on *F* expression could then allow for local evolution of a female heterogamety system.

To test this reasoning, we used simulations in which we varied two parameters: the rate *β* at which temperature increases *F* expression (see Equation 1) and the threshold value *θ*_F_ for feminization. Here, we exclude Y-chromosomal fitness effects, but explore their impact on SD evolution in the following section. We found that *F*_I_ can spread in the entire metapopulation when the feminization threshold *θ*_F_ is sufficiently small (Figure 2A), regardless of how strongly temperature affects *F* expression. For sufficiently high values of *θ*_F_, *F*_I_ was unable to spread in colder demes because it would result in intersexual development (Supplementary Figure 4), but could still spread in warmer demes. Under these conditions, a geographical cline in the frequency of *F*_I_ evolves (Figure 2B).

**Figure 2:**
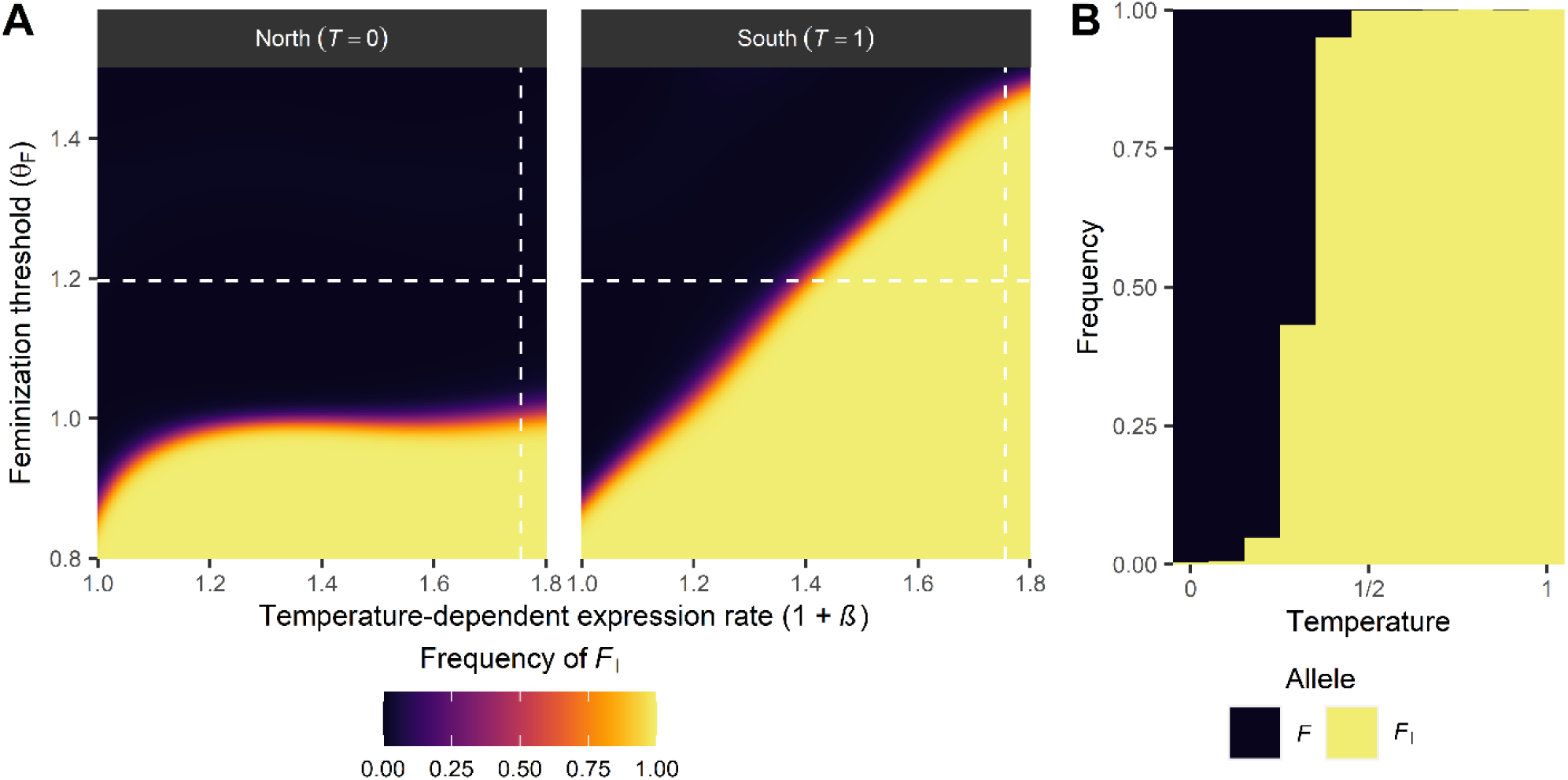
Conditions for spread of a dominant female-determining gene *F*_I_ as a function of the temperature-dependent expression rate 1 + *β* and the feminization threshold *θ*_F_. When *β* = 0, expression is unaffected by temperature. (A) Equilibrium frequencies of *F*_I_ at edges (first/last deme) of the population range; local temperatures are indicated in brackets (parameter values: *θ*_*M*_ = 0.2; *µ*_D_ = 0.001). In the northern deme (*T* = 0), temperature-dependent expression of *F* is lowest while expression is highest in the southern deme (*T* = 1). (B) Between these two extremes, the equilibrium frequency of *F*_I_ increases along the temperature gradient (shown here for *θ*_F_ = 1.20; *θ*_M_ = 0.2; *β* = 0.76; *µ*_D_ = 0.014; the results depicted were chosen based on whether or not a gradient in *F*_I_ was observed, with parameter values in each simulation being chosen at random from uniform distributions). Depicted in A and B are the frequency of the *F*_I_ allele at the maternally-inherited locus in females. White dashed lines in A indicate the parameter values for the simulation results depicted in B (vertical line: 1 + *β*; horizontal line: *θ*_F_).

These results underline that the distribution of *F*_I_ is constrained by the expression level of *F*, and show that temperature-dependent effects on gene expression can establish gradients when temperature varies. Due to its feminizing effect, an *F*_I_ allele is transmitted as if it were a female-limited W-chromosome in a ZW system, wherein males have a ZZ genotype, whereas females have a ZW genotype (in contrast to XY systems where females are XX and males are XY). As a result, it occurs only in *F*_I_/*F* females (or non-reproducing intersexes). In the presence of *M* product, the *F*_I_ product is not broken down but the product of regular sensitive *F* alleles is degraded, so that regular *F* alleles do not contribute to the total *F* product. This scenario becomes increasingly probable as either *M*_Y_ and/or *M*_A_ frequency increases, as more *F*_I_-bearing individuals will also bear *M*_Y_ and/or *M*_A_. Therefore, feminization of developing embryos under these conditions is achieved solely by the activity of the *F*_I_ allele. Because *F*_I_ is insensitive to *M*, the total *F* activity is determined by its expression (see Equation 1). When no temperature-dependent expression occurs, feminization is only achieved when the baseline genetic expression level exceeds the feminization threshold, but in the presence of temperature effects is less constrained. Therefore, when *θ*_F_ is sufficiently low, *F*_I_ can spread everywhere independent of temperature (Figure 2A, left panel), whereas otherwise *F*_I_ frequencies depend on the temperature-dependent expression rate (Figure 2A, right panel) and the local temperature (Figure 2B).

### Y-chromosomal fitness effects modulate the conditions under which SD turnover can occur and can enable complex SD polymorphisms

For the simulations discussed above, we assumed that *M*_Y_ is not associated with any fitness effects. Under these conditions, *M*_Y_ was always lost and replaced by *M*_A_ because *M*_Y_ instead was neither favoured by selection nor recurrently formed through mutation. However, as *M*_Y_ is the ancestral SD gene, it may have induced the chromosome on which it is located to undergo Y-chromosome evolution (reviewed in ^29,30^). If so, this chromosome is expected to become enriched for sexually antagonistic (SA) genetic variants as well as recessive deleterious mutations. In effect, this would cause *M*_Y_ to be associated with higher fitness in heterozygous *+/M*_Y_ males, but to induce a fitness cost to *M*_Y_-bearing females (who also carry *F*_I_) as well as all homozygous *M*_Y_/*M*_Y_ individuals. Both sexually antagonistic genetic variation and a cost of homozygosity can in theory strongly affect evolutionary transitions in SD (reviewed in ^31^). We therefore performed additional simulations where we considered a sexually antagonistic fitness effect of the Y-chromosome (see Supplementary Table 1), causing the Y-chromosome to affect individual fitness during the mating phase or during development (positively in males, negatively in females). In addition, we include costs of *M*_Y_/*M*_Y_ homozygosity as a form of viability selection during embryogenesis.

When *M*_Y_ is under sexually antagonistic selection, we find that it is maintained over *M*_A_ (Figure 3). Sexually antagonistic selection on *M*_Y_ can also inhibit invasion of *F*_I_ if selection is sufficiently strong, so that negative effects of *M*_Y_ in females reduces the fitness of *F*_I_/*F* females relative to non-*M*_Y_-carrying *F/F* females. However, when the rate of introduction of novel *M*_A_ alleles is sufficiently high, *F*_I_ can always invade even if the selective effects associated with *M*_Y_ are strong. When *M*_A_ originates more frequently and therefore reaches higher frequencies, a more male-biased sex ratio results (similar to Y-chromosomal meiotic drive^32,33^) and hence the selective benefit for *F*_I_ as a female-determining factor increases.

**Figure 3:**
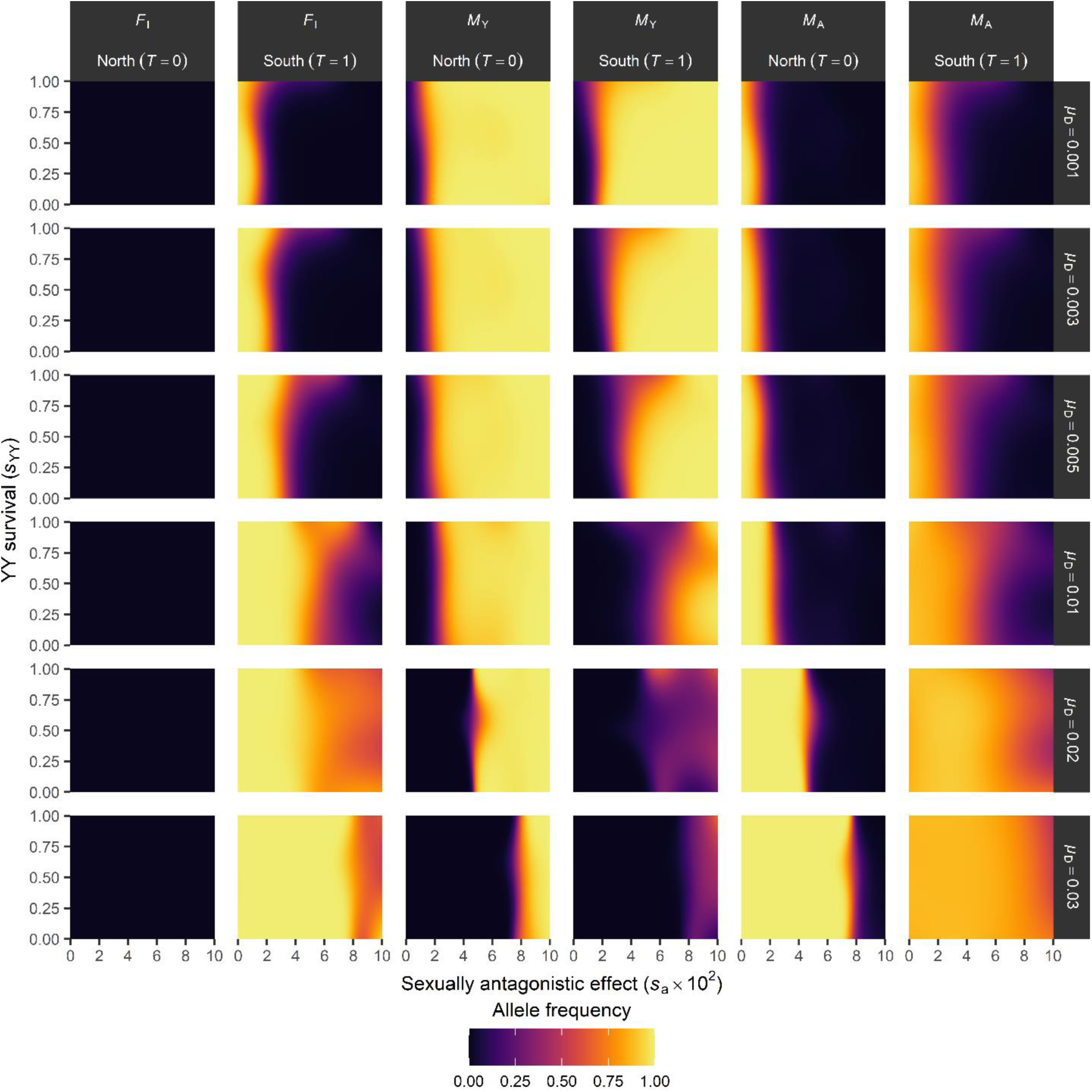
Y-chromosomal fitness effects alter the scope for SD transitions. Shown are the predicted equilibrium frequencies of *F*_I_ in females (maternally-inherited alleles), *M*_Y_ and *M*_A_ in males (paternally-inherited alleles); we restrict our analysis to the maternal (*F*_I_) c.q. paternal (*M*_Y_, *M*_A_) alleles to account for their (potential) sex-specific transmission. Horizontal labels indicate locus and temperature, vertical labels the *M*_A_ activation rate *µ*_D_. Further parameter values used in simulations: *θ*_F_ = 1.2; *θ*_M_ = 0.3; *β*_T_ = 0.5. Predicted allele frequencies were smoothed with binomial generalized additive models.

Y-chromosomal fitness effects can also enable maintenance of both *M*_Y_ and *M*_A_ in the population, albeit in different locations. This occurs when *M*_Y_ is disfavoured through reduced survival of YY homozygotes in subpopulations with *F*_I_, in contrast to subpopulations without *F*_I_, where *M*_Y_ is favoured over *M*_A_ via sexually antagonistic selection in +/*M*_Y_ heterozygotes. Such *M*_Y_ *versus M*_A_ polymorphism is highly similar to the distribution of Y-chromosomal *versus* autosomal *M*-factors in the housefly *M. domestica*. In this species, autosomal *M*-factors are more prevalent at lower latitudes and coincide with a dominant feminizing allele *tra*^D^ (which is equivalent to *F*_I_ as described above), resulting in three latitudinal gradients in Y-chromosomal *M*-factors, autosomal *M*-factors, and the *tra*^D^ allele. We find that Y-chromosomal fitness effects enable the evolution of this complex system in our model (see Box 1). This provides an adaptive explanation for the evolution of this system in contrast to existing models of SD evolution, which have been unable to predict when stable polymorphisms for *tra versus tra*^D^ along with Y-chromosomal *M*-factors *versus* autosomal *M*-factors may occur in general, and in particular when these coincide along environmental clines as observed under natural conditions.

## Discussion

We have presented a model for the evolution of SD systems in a context where sex is determined by genetic factors in combination with environmental effects. In our model, female development is induced when the activity of a feminizing gene *F* exceeds a certain threshold, whereas male development is achieved when *F* activity is below a different and lower threshold. This can be caused by inhibition of *F* activity by the maleness-promoting gene(s) *M*_Y_ and/or *M*_A_, or by loss of *F* expression. We incorporated a positive effect of temperature on the expression of a feminizing locus *F*. We find that several different SD systems can be realized depending on the activity of an *F* allele relative to the threshold values for masculinization and feminization. Temperature-dependent effects on *F* expression may alter the relationships between *F* activity and SD thresholds, thereby enabling the evolution of different genetic SD systems. A particular prediction is the transition from male heterogamety to female heterogamety, which occurs in our model via the evolution of an insensitive dominant feminizing variant (*F*_I_) that induces femaleness even in the presence of *M*_Y_ and/or *M*_A_. *F*_I_ can spread when activity of a single *F* or *F*_I_ allele is sufficient to induce feminization; when activity is modulated by temperature, this can lead to local invasions of *F*_I_ and subsequently differentiation between populations along temperature gradients. Differentiation can also occur for *M*_Y_ and *M*_A_, with *M*_Y_ being favoured in absence of *F*_I_ and *M*_A_ in presence of *F*_I_. This occurs when *M*_Y_ is associated with certain fitness effects such due to linkage with sexually antagonistic variants or recessive deleterious mutations. Altogether, we show that this can lead to the coexistence of multiple gradients in SD genes as found in *M. domestica*.

Our model can be amended to other SD systems than the *M. domestica* system on which it is based, provided that they have a basic GSD framework influenced by an environmental effect. Environmental effects on genetic sex determination systems are being reported in an increasing number of species^16,34^. Although temperature-dependent effects are well-documented, other environmental effects may also influence SD in certain systems such as hormonal imbalance in fish due to pollution^4^. Other components and assumptions of the model that are based on the housefly system may be represented differently in other species but with similar effects. For example, the impact of *M*_A_ evolution is not due to a specific mechanism of mutation, but more generally by causing a male-biased sex ratio, thereby promoting the invasion of *F*_I_. Male-biased sex ratios occur due to *M*_A_ overrepresentation in the gene pool via its *de novo* evolution, but the same effect can be achieved by translocation of a Y-chromosomal male-determining gene or via association with meiotic drivers^33,35^. Additionally, we see that in absence of an association between *M*_Y_ and fitness effects, *M*_A_ replaces *M*_Y_ altogether, yet still drives the invasion of *F*_I_, showing that our model does not strictly require a third locus. Inversely, it is likely that a more complex genetic basis generates similar evolutionary patterns, such as when various genes can evolve into a male-determining gene^28^. In this scenario, many genes having a small chance to evolve into a male-determining gene may have the same net effect on sex ratio as a gene that is prone to evolving a masculinizing function. In this light, it will be interesting to test whether genes that have acquired ex-determining function in one species are prone to evolve a similar function in a related species, where it has no sex-determining function.

Previous work has shed light on the evolution of ESD and GSD systems, and when transitions between these two may arise^10,20,36^. However, our understanding of the evolution of polymorphic SD systems and the potential for environmental heterogeneity to influence this process remains limited. Our results highlight environmental effects on GSD systems, and under which conditions this can lead to within-species polymorphism in GSD systems. Moreover, even in systems that appear to be fully GSD, the role of environmental influences on the SD processes must not be ignored as these may have played an important role in their evolution. In extension of this, changes in environment, e.g., due to global warming, may impose yet unforeseen selective pressures on species with GSD systems.

### Box 1: Evolution of the polymorphic housefly system

Our model has been inspired by the common housefly *Musca domestica*, in which wild populations harbour different SD systems (reviewed in ^14^). Here, we discuss how our model can explain the evolution of this system.

In houseflies, sex is determined by a linear cascade of genes. First, *transformer* (*tra*) induces female development when active in developing embryos^37^; its activity depends on pre-mRNA splicing, which is sensitive to temperature as well as other stressors^38^. Second, masculinizing factors (*M*-factors) such as *Mdmd* ^39^ trigger male development by inhibition of the *tra* loop through splicing of *tra* pre-mRNA to a male-specific variant. Intriguingly, the *M*-factors in *M. domestica* are found on different chromosomes in different populations. There also exists an insensitive variant of *tra, tra*^D^, that induces female development in all carriers regardless of whether they carry any *M*-factors. In our model, *F* represents *tra* and likewise *tra*^D^ corresponds to the dominant *F*_I_ as discussed in the main text; *M*_Y_ and *M*_A_ in our model represent *M*-factors.

High-latitude *M. domestica* populations have a male heterogamety (XY) system in which the Y-chromosomal *Mdmd* gene induces maleness, and all individuals carry two regularly-sensitive *tra* alleles. At lower latitudes, however, females usually carry the insensitive *tra*^D^ allele, and both sexes can be homozygous for an autosomal copy of *Mdmd*; hence, these populations have a female heterogamety (ZW) system (Figure 4; ^14,40^). The geographical transition between these two SD systems is gradual, so that clines exist in the frequencies of Y-chromosomal *Mdmd* (decreasing towards lower latitudes), autosomal *Mdmd* and *tra*^D^ (both increasing towards lower latitudes). Temperature likely plays a causal role in maintaining these gradients by affecting the SD process^41–44^. Temperature effects on housefly SD have been reported in the form of biased sex ratios produced in wildtype crosses^42^ as well as in females carrying the *masculinizer* (*man*) mutation, another variant of *tra*^21,37^. The *man* mutation represents a maternal-effect male-determining gene, where *man*-carrying females can produce all-male progeny even if the progeny do not carry an *M*-factor. However, this effect is incomplete and temperature-sensitive^21^, with offspring sex ratios more male-biased at higher temperatures. Altogether, temperature seems to have an important influence on SD in *M. domestica*, but the underlying mechanisms are not yet fully understood.

**Figure 4:**
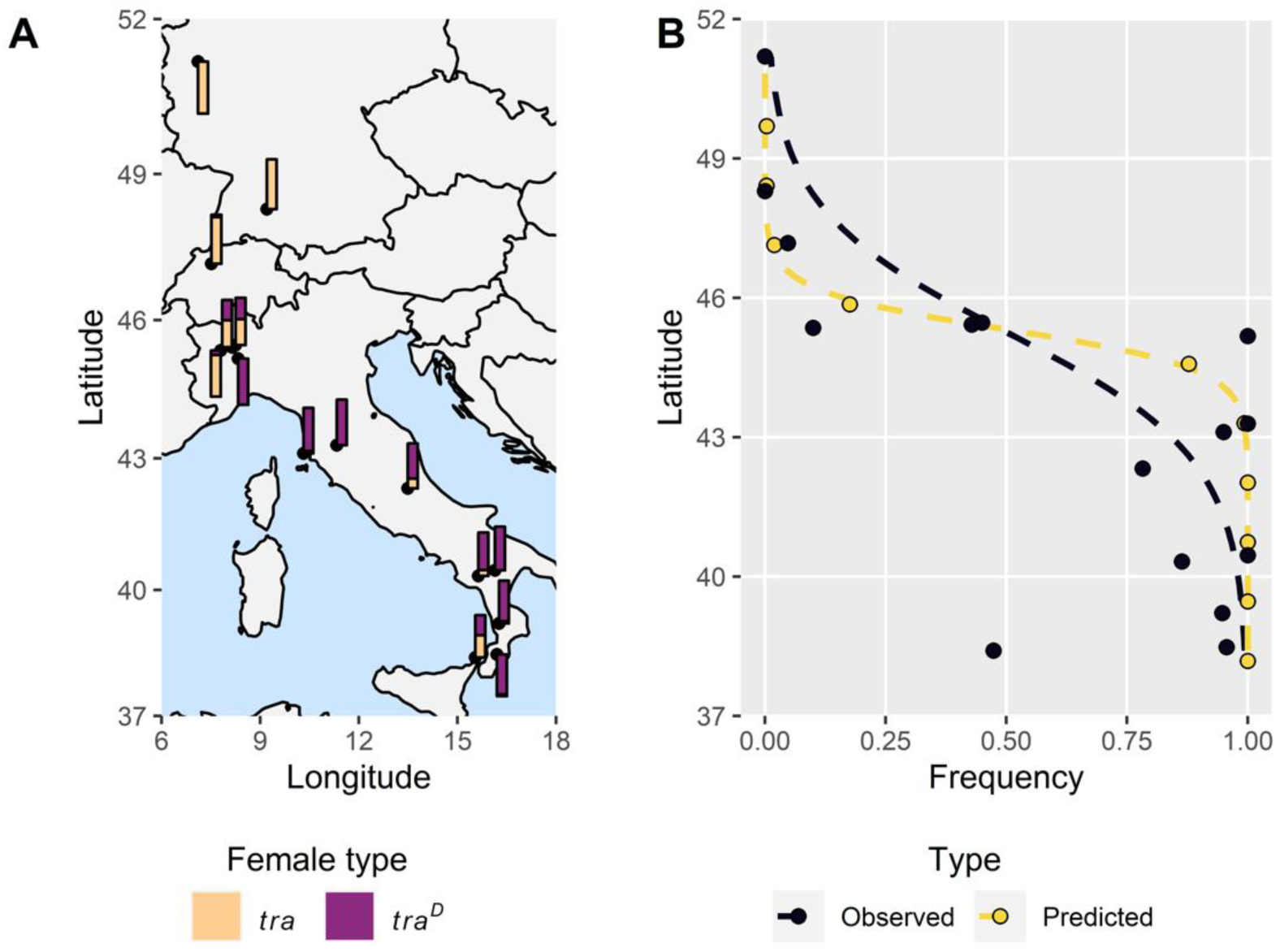
Model predictions vs. the observed latitudinal frequency gradient of a female-determining gene in the housefly *Musca domestica*. (A) The frequency of *tra/tra* (light orange) and *tra*^D/^/*tra* (purple) females in Europe. Adapted from ^40^. (B) Observed (black) and predicted (yellow) frequencies of *tra*^D^-bearing females (deme position adjusted to match observed latitude range). Dashed lines indicate fitted binomial GLMs for allele frequency with latitude as the sole predictor variable. Parameter values used: *β* = 0.5; *θ*_F_ = 1.15; *µ*_D_ = 0.005; *s*_*a*_ = 0.05; *s*_*YY*_ = 0.9; *d* = 0.1.

The housefly with its different SD systems is represented in our model by gradients in the frequencies of *M*_Y_ (decreasing with temperature), *M*_A_, and *F*_I_ (both increasing with temperature). Presumably, *tra*^D^ is limited to warmer localities for similar reasons as *F*_I_ in our model, i.e., because a single *tra*^D^ allele may not be sufficient to induce feminization at low temperatures. Y-chromosomal *Mdmd* and autosomal *Mdmd* may follow similar dynamics as *M*_Y_ and *M*_A_. Y-chromosomal fitness effects can yield gradients in *M*_Y_ and *M*_A_, but may also prevent the spread of *F*_I_, particularly when novel *M*_A_ alleles enter the population at a low rate. The evolution of a housefly-like polymorphic SD system therefore depends on a balance between the Y-chromosomal fitness effects and the rate at which new autosomal *Mdmd* arises. In our model, we find that a housefly-like system can evolve under various rates of *M*_A_ *de novo* evolution (Figure 5). Higher rates of *M*_A_ evolution require stronger SA effects for *M*_Y_ to be maintained in low-temperature demes. Costs of YY homozygosity do not appear essential for *M*_Y_ to be lost in the presence of *F*_I_, although they may increase the likelihood of *M*_Y_ being lost in favour of *M*_A_ in the presence of *F*_I_ by reducing the fitness of YY homozygotes. Possibly, costs of *M*_Y_ in females due to sexually antagonistic selection may suffice to promote the loss of *M*_Y_. Our model therefore can explain the evolution of complex SD systems such as those found in houseflies. Thus, we propose a novel hypothesis for the evolution of the housefly system in which the sex determination cascade is shaped by a combination of environmental influences on *tra*, recurrent evolution of autosomal *Mdmd*, and fitness effects associated with the Y-chromosomal copy of *Mdmd* (Figure 6).

**Figure 5:**
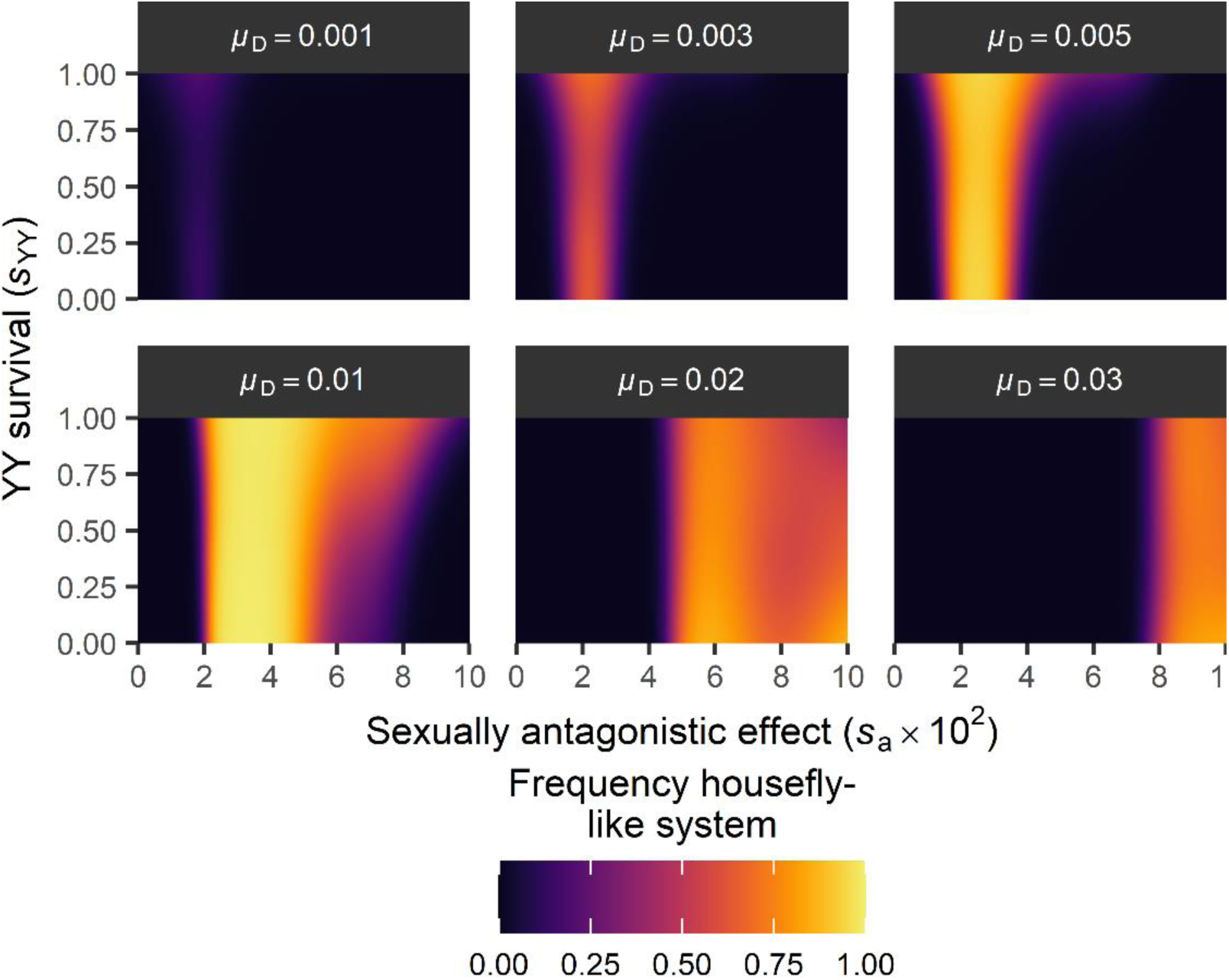
Evolution of a housefly-like SD system. A housefly-like SD system is defined by *M*_Y_ being the major allele at *T* = 0 but the minor allele at *T* = 1 (in males, paternally-inherited allele), and vice versa for *M*_A_ (in males, paternally-inherited alleles) and *F*_I_ (in females, maternally-inherited alleles). Frequency denotes the predicted frequency of observing a housefly-like system at equilibrium in the model. Parameter values: *θ*_F_ = 1.2; *θ*_M_ = 0.3; *β* = 0.5. Simulations were scored as exhibiting a housefly-like system as described above (10000 simulations in total). To obtain these predicted scores, we fitted a binomial generalized additive model.

**Figure 6:**
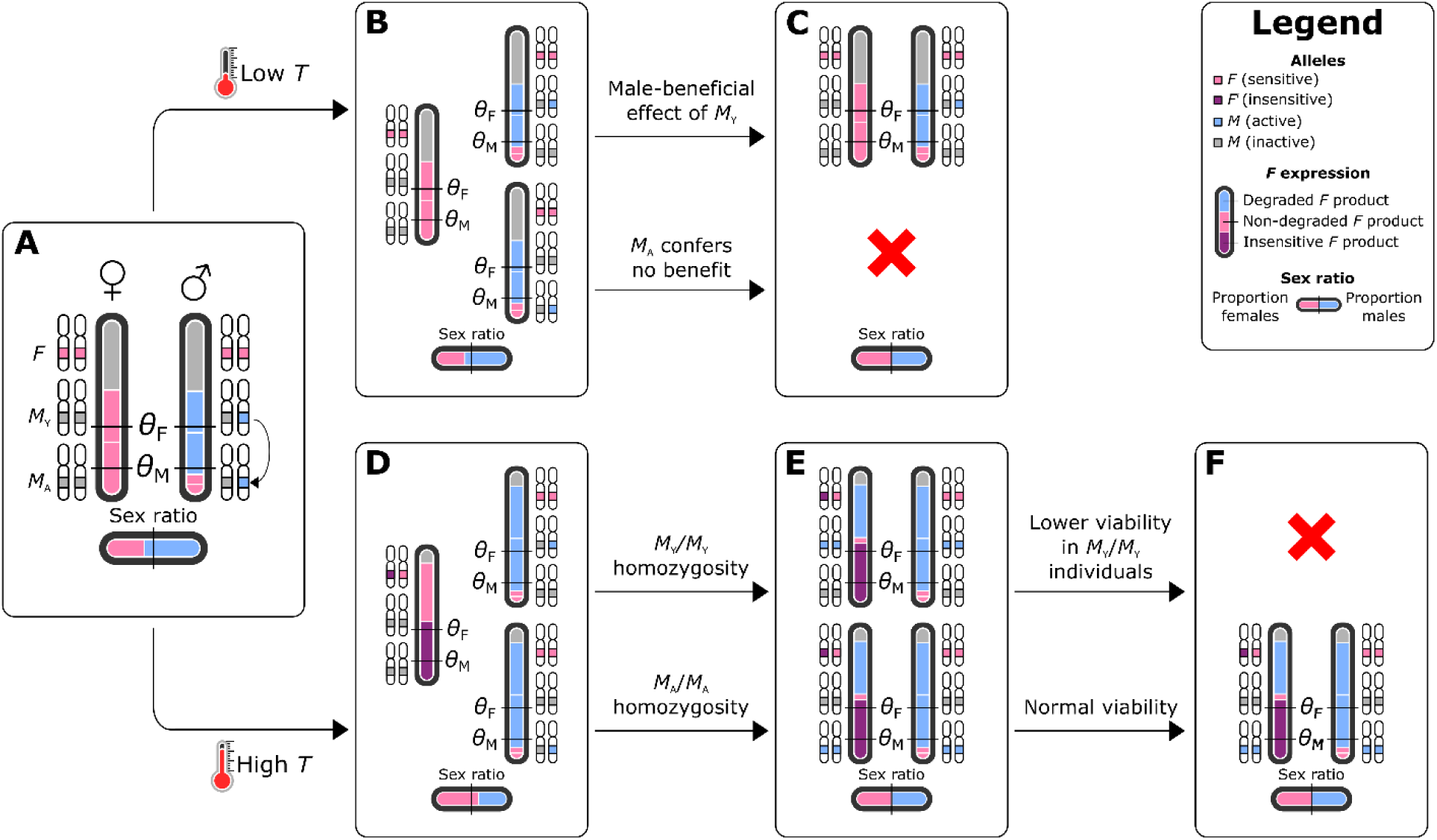
A novel hypothesis for the evolution of the housefly polymorphic sex determination mechanism. (A) Evolution of *M*_A_ (here represented by transposition of *M*_Y_) results in an excess of *M* factors in the population and a male-biased sex ratio. (B) At low temperatures, the *M*-insensitive *F*_I_ allele (equivalent to *tra*^D^ in *M. domestica*) cannot evolve, and both *M*_Y_ and *M*_A_ persist in a heterozygous state. (C) Because *M*_Y_ is associated with a fitness benefit in males, *M*_Y_-bearing males outcompete *M*_A_-bearing males, resulting in a return to the ancestral state with *M*_Y_ as the sole male-determining allele in a XY system. (D) In contrast, at high temperatures, the *F*^D^ allele can evolve, and has a fitness benefit as a result of sex ratio selection. (E) As *F*^*D*^ spreads, *M* alleles can be transmitted by females resulting in the formation of homozygous *M*_Y_//*M*_Y_ and *M*_A_//*M*_A_ individuals. (F) Because *M*_Y_ homozygosity is associated with a viability cost, these individuals have lower fitness than *M*_A_/*M*_**A**_ individuals. This results in a loss of the *M*_Y_ allele and fixation of the *M*_A_ allele in its place as a co-factor for male development. In effect, a transition has occurred from XY male heterogamety to ZW female heterogamety as the sex-determining role has been taken over by *F*^D^.

## Acknowledgements

We thank the Center for Information Technology of the University of Groningen for providing access to the Peregrine high-performance computing cluster; Pina Brinker, Fangying Chen, and Peter Hoitinga for feedback on an earlier version of the manuscript; and Babak Arani and the Evolutionary Genetics cluster for fruitful discussion. MAS was supported by an Adaptive Life grant awarded to LWB, IP, and J-CB by the University of Groningen.

## Data availability

Source code and analysis scripts for all simulations are freely available via GitHub (https://github.com/MartijnSchenkel/Environmental_GSD_evolution).

## Methods

An overview of all parameters and their default values are provided in Supplementary Table 2.

### Life cycle

We simulate a population consisting of individuals distributed among *N* demes (*K* individuals per deme, for a total population size *NK*) arranged along a linear gradient (Figure 1B); we vary the environmental cue *T* normalized from 0 in the first deme to 1 in the last deme. *T* positively affects the expression of a feminizing locus *F* and may thereby increase the probability of an individual developing as a female. In each deme, we generate a fixed number of *K* individuals upon initiation. Individuals have a diploid genome consisting of three linkage groups, of which one carries the *F* locus. The two remaining linkage groups can carry a male-determining *M* locus, whose product *M* degrades the *F* product. We designate one of these linkage groups as being the original sex chromosome pair and accordingly refer to these as XY chromosome pair; we refer to the *M*-locus on this chromosome pair as *M*_Y_. The other linkage group is referred to as the autosomal chromosome pair, and we refer to its *M*-locus as *M*_A_. Sex is determined by the total amount of *F* product available after interacting with *M*. We assume a male heterogametic system in which females carry two X-chromosomes that lack an active *M* allele, and males carry one X-chromosome and a Y-chromosome that harbours an active *M*_Y_ allele. All individuals initially lack active *M*_A_ alleles. Reproduction occurs by random mating between males and females within each deme, during which all linkage groups follow Mendelian segregation. Each allelic trait can mutate with a certain trait-specific probability. Reproduction results in a total of *K* juveniles within each deme, which may then disperse with a probability *d* to a random neighbouring deme (or *d*/2 in the first/last demes). After dispersal, all adults die and are replaced by the juveniles. For all simulations, a “burn-in” period of 20,000 generations is applied during which all demes have a *T* = 0. After this, we increase *T* in each deme to its final value (as determined by their position in the gradient) during a warmup period of 10,000 generations. We use a burn-in and warmup phase to ensure that the system can evolve to a stable state prior to incorporating environmental effects and to ensure selection due to environmental effects does not change abruptly. After the warmup phase, we keep conditions stable for 200,000 generations to allow the system to evolve to a new equilibrium. We then analyse the genotypes of all individuals to determine which SD systems have evolved in which demes.

### Sex determination

An individual’s sex is determined by the net activity *z*_*F*_ of the two alleles at the *F* locus, based on the initial expression level of each *F* allele minus the amount of *F* product that is degraded by *M* (see Figure 1A). Initially, each allele produces an amount 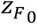 of F product which has a sensitivity 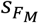 to breakdown by *M*. Environmental effects on SD are included solely with regard to *F* expression, so that the gross expression 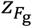 (prior to eventual breakdown by *M*) of an F allele is given by:

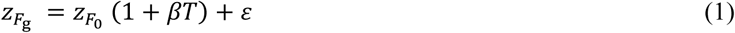

Here, *β* refers to the rate at which *F* expression increases between *T* = 0 and *T* = 1. To model within-deme heterogeneity in *T* and developmental noise, we add some noise to the expression of *F* by adding a Gaussian amount *ε* with *µ* = 0 and *σ* = *σ*_*F*_. The cumulative amount 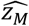 of *M* product is determined by summing up the expression levels *z*_*M*_ of all *M* alleles. *F* breakdown by *M* is determined per *F* allele by the expression level and sensitivity of the product, so that the net activity *z*_*F*_ of an *F* allele is given by:

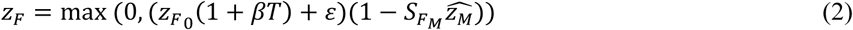

We sum up the net activity of both alleles of *F* to obtain the net activity of the *F* locus, 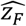, based on which sex is determined:

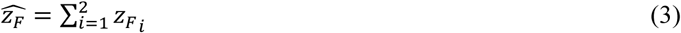

in which *i* is used to indicate the maternal and paternal allele. Individuals develop into males if 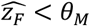 or into females if 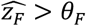. If 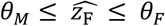 individuals develop into infertile intersexes. *F* expression and sensitivity, as well as *M* expression, can vary from 0 to 1.

### Reproduction and mutation

Reproduction occurs by mating between females and males residing in the same deme; an individual’s probability of being sampled as a mate is proportional to its fitness relative to other same-sex individuals and depends solely on sex chromosome genotype; in absence of Y-chromosomal fitness effects mating effectively occurs between randomly-sampled males and females. Mutations can occur within each gamete, and occur independently for *F* sensitivity, *F* expression, and *M* expression each with a trait-specific probability *µ*. Mutations may either result in a certain shift in trait value (regular mutations) or in a loss-of-function type mutation where the trait value is set to zero (null mutation). When mutations occur, they have a trait-specific probability *µ*_null_ of being a null mutation. Regular mutations result in a change in a trait value by summing the current trait value with a value sampled from normal distributions with *µ* = 0 and standard deviations 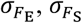, and 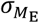 for *F* sensitivity, *F* expression, and *M* expression. Once a trait value equals zero through either regular or null mutations, we consider this to be a loss of function and prevent the trait from undergoing further mutation so that there is no gain of function. New (expressed) *M*_A_ alleles may arise *de novo* with a frequency *µ*_D_. The expression level of newly-evolved *M*_A_ locus is sampled from a normal distribution with mean *µ*_E_ and standard deviation *σ*_E_.

### Fitness effects of the Y-chromosome

We incorporate two possible fitness effects that are associated with *M*_Y_. First, we incorporate a sexually antagonistic fitness effect of the Y-chromosome with additive fitness effects −*s*_a_ and *s*_a_ in females and males (see Supplementary Table 1). This results in a positive effect in males but a negative effect in females, and resembles a scenario in which the Y-chromosomal *M*-locus is tightly linked to one or more sexually antagonistic alleles. These fitness effects are effectuated during the mating phase of our model. Second, we incorporate a fully recessive viability cost of *M*_Y_/*M*_Y_ homozygosity, whereby developing YY individuals survive to the adult stage with a probability *s*_YY_ ≤ 1. This resembles the effect of mutation accumulation on the Y-chromosome. For both fitness effects, we assume that *M*_Y_ is fully linked to the loci under selection so that no recombination occurs.

### Data analysis

We categorize all *F* alleles based on their sensitivity level and their expression level, where an *F* allele is considered insensitive if its sensitivity is equal to 0 (and sensitive if it is larger than 0), and is unexpressed when its expression is less than *θ*_M_/2 (and expressed if otherwise). We classify all alleles that are insensitive and expressed as *F*_I_ alleles, and all others as *F* alleles. We categorize all *M* alleles based on their expression level; we consider an *M* allele unexpressed when its expression is lower than (2 − *θ*_F_)/2, and expressed if its expression equals or exceeds (2 − *θ*_F_)/2. The threshold for *F* expression is based on the assumption that an expression value below *θ*_M_/2 is insufficient to prevent maleness when an individual is homozygous. Similarly, an *M* allele may break down a small amount of *F* product without preventing female development. Assuming *F* is fully expressed (and hence a total of 2 *F* product is generated), breakdown by M can at most be 2 − *θ*_F_. When both *F* alleles are fully sensitive, this means that an individual can be homozygous for an M that breaks down at most (2 − *θ*_F_)/2 and still develop into a female.

Following categorization, for each deme we recorded the frequencies of *F* and *F*_I_ as well as unexpressed and expressed *M* alleles on the maternally-inherited and paternally-inherited alleles. We used the ‘mgcv’ package^45^ to fit generalized additive models (GAMs) with binomial distributions on the allele frequencies at the two extremes *T* = 0 and *T* = 1 for each locus. For the *F* locus, we did so on the allele frequency of the *F*_I_ allele on the maternally-inherited copy and for both *M*_Y_ and *M*_A_ we fit GAMs on the frequency of expressed alleles on the paternally-inherited copies. In each GAM, we included a full tensor product smooth between the parameters of interest.

To assess whether a polymorphic system resembling that found in natural housefly populations had evolved, we determined for each simulation whether (1) *F*_I_ was the minor allele at *T* = 0 and the major allele at *T* = 1; (2) *M*_Y_ was the major allele at *T* = 0 and the minor allele at *T* = 1; and (3) *M*_A_ was the minor allele at *T* = 0 and the major allele at *T* = 1. Minor (major) alleles are defined as having a frequency below (over) ½. Simulations that met all three criteria were considered to have evolved a housefly-like SD system. The resulting scores were used to fit a binomially-distributed GAM with a full tensor product smooth between *M*_A_ de novo rate (*µ*_D_), sexually-antagonistic fitness effect (*s*_*a*_), and YY survival rates (*s*_YY_).

All data analysis was carried out in R (v.4.0.0; ^46^) and RStudio (v.1.2.5033; ^47^), using the ‘cowplot’^48^, ‘maps’^49^, ‘mapsproj’^50^, ‘mgcv’^45^, ‘tidyverse’^51^, and ‘viridis’^52^ packages.

## Authorship contributions

LWB and IP conceived the study; MAS, J-CB, LWB and IP designed the model; MAS and IP generated the model source code; MAS conducted the simulations, performed the data analysis, wrote the initial draft; MAS, J-CB, LWB and IP contributed to writing the final manuscript.

## Supplementary Material

### Appendix 1: F activity relative to SD thresholds determines which SD systems are evolutionary stable

As an initial exploration of our model, we performed a set of 10,000 simulations without temperature-dependent overexpression, but for different values for the SD thresholds *θ*_M_ and *θ*_F_. In addition, for these simulations we did not incorporate fitness effects associated with *M*_Y_. For each simulation, we determined the most prevalent genotype among females and males at equilibrium, based on the expression and sensitivity levels of their *F* alleles as well as the number of expressed *M*_Y_ and *M*_A_ alleles. We categorized simulations according to the activity of a single *F* allele (*z*_*F*_) relative to *θ*_M_ and *θ*_F_ to determine which SD systems can evolve under different levels of *F* activity.

We find that under different relationships between the maximum potential activity of a single *F* allele (*z*_*F*_) and the SD thresholds, different SD systems can evolve (Table 1). When *θ*_M_ < *z*_*F*_ < *θ*_F_, nearly all simulations yield an SD system where both sexes have two sensitive and expressed *F* alleles, and males are heterozygous for a single *M* (either *M*_Y_ or *M*_A_; females *F/F; +/+*, males *F/F; +/M*, with + indicating the absence of an expressed *M* allele). In contrast, when *z*_*F*_ < *θ*_M_ < *θ*_F_ or *θ*_M_ < *θ*_F_ < *z*_*F*_, we additionally encounter systems in which *F* alleles evolved to become insensitive and/or unexpressed, but the distribution of these alleles over the sexes differs between these two scenarios: when *z*_*F*_ < *θ*_M_ < *θ*_F_, females carry two insensitive F alleles, whereas males carry a single insensitive F and one unexpressed F; here, insensitive *F* alleles can be regarded as recessive female-determining alleles, whereas the sensitive and expressed *F* allele (in presence of expressed *M*) or unexpressed *F* plays the role of a dominant male-determining allele. In contrast, when *θ*_M_ < *θ*_F_ < *z*_*F*_, females may carry a single insensitive *F* allele, whereas males carry none, suggesting that insensitive *F* alleles are dominant female-determining alleles. This is corroborated by the presence of expressed *M* alleles in both sexes in the simulations, which would otherwise induce maleness in their carriers.

In simulations where insensitive *F* alleles have evolved, we find that the remaining *F* alleles have become unexpressed, and in some cases that expressed *M* alleles are lost as well. In these simulations, insensitive *F* alleles spread first, along with an increased frequency of expressed *M* alleles in both sexes (Supplementary Figure 1), the latter being required to achieve an equal sex ratio. Next, we see that sensitive *F* alleles become unexpressed (and subsequently insensitive due to the constant mutation pressure affecting *F* sensitivity); the insensitive expressed *F* allele is retained as it now performs the SD function. Along with the increase in unexpressed *F* alleles, we similarly find that M alleles lose their expression. Based on these dynamics, we infer that evolution of *F* insensitivity indirectly renders the loss of expression selectively neutral for sensitive *F* alleles, which in turn renders the loss of *M* expression neutral; both genes may therefore decay via mutation accumulation in a stepwise order (Supplementary Figure 3). In addition to the systems identified in our simulations, we speculate some other systems may also be evolutionary stable (Supplementary Figure 2). Their absence from our simulations may be because they only rarely arise through evolution, or because they represent intermediate states between some of the systems that are observed. The latter for example applies to systems where females have one insensitive and one sensitive F allele, males have two sensitive F alleles, and *M* is fixed in both sexes (females *F*_I_*/F; M/M*, males *F/F; M/M* in Figure 3). This system is prevalent in some simulations prior to the accumulation of unexpressed F alleles and loss of M (e.g., Supplementary Figure 1). In absence of fitness effects related to *M*_Y_, recurrent *de novo* mutation of *M*_A_ results in *M*_A_ replacing *M*_Y_ as the male-determining factor; without evolution of *F*, this represents a transition from one male heterogamety system to another, as *M*_Y_ and *M*_A_ perform equivalent functions. Male heterogamety systems with *M*_Y_ or *M*_A_ as the male-determining gene are observed regardless of the activity of *F* relative to the SD thresholds (Supplementary Table 3). Moreover, male heterogamety with *M*_Y_ or *M*_A_ as a dominant male-determiner is observed in all simulations when *θ*_M_ < *z*_*F*_ < *θ*_F_.

## Supplementary Tables

**Supplementary Table 1:** Sexually antagonistic fitness effects of *M*_Y_. *s*_*a*_ denotes the sexually antagonistic fitness effect of *M*_Y_ in females and males, whereas *h*_F_ and *h*_M_ denote the dominance of these fitness effects in *+/M*_Y_ heterozygotes. We assume *M*_Y_ to be deleterious to females but beneficial to males.

| | $+/+$ | $+/M_Y$ | $M_Y/M_Y$ |
| --- | --- | --- | --- |
| Female | 1 | $1 - h_F s_a$ | $1 - s_a$ |
| Male | 1 | $1 + h_M s_a$ | $1 + s_a$ |

**Supplementary Table 2:** Parameters in the model and their default values.

| Variable | Description | Standard value |
| --- | --- | --- |
| $N$ | Number of demes | 11 |
| $K$ | Population size per deme | 10,000 |
| $d$ | Probability of dispersal by an individual after maturation | 0.1 |
| $\theta_F$ | Lower threshold for female development | 1.2 |
| $\theta_M$ | Upper threshold for male development | 0.3 |
| $z_{F0}$ | Baseline expression level of $F$ alleles | 0.7 |
| $S_{FM}$ | Sensitivity of $F$ allele's product to breakdown by $M$ | 0.95 |
| $z_{M_Y}$ | Expression level of $M_Y$ alleles | 0.9 |
| $z_{M_A}$ | Expression level of $M_A$ alleles | 0.9 |
| $s_{YY}$ | Survival rate of $YY$ individuals | 1 |
| $h_F$ | Dominance of $M_Y$ sexually antagonistic fitness effect in females | 0.5 |
| $h_M$ | Dominance of $M_Y$ sexually antagonistic fitness effect in males | 0.5 |
| $s_a$ | Sexually antagonistic effect of $M_Y$ (negative in females, positive in males) | 0 |
| $\mu_{\text{null}}$ | Proportion of mutations that are null mutations | 0.01 |
| $\mu_E$ | Mutation rate for $F$ expression level | 0.1 |
| $\sigma_{FE}$ | SD for mutation effect for $F$ expression level | 0.1 |
| $\mu_{E_{\text{null}}}$ | Proportion of mutations in $F$ expression that are null mutations | $\mu_{\text{null}}$ |
| $\mu_S$ | Mutation rate for $F$ sensitivity | 0.1 |
| $\sigma_{F_S}$ | SD for mutation effect for $F$ sensitivity | 0.1 |
| $\mu_{S_{\text{null}}}$ | Proportion of mutations in $F$ sensitivity that are null mutations | $\mu_{\text{null}}$ |
| $\mu_M$ | Mutation rate for $M$ expression level | 0.1 |
| $\sigma_{M_E}$ | SD for mutation effect for $M$ expression | 0.1 |
| $\mu_{M_{\text{null}}}$ | Proportion of mutations in $M$ expression that are null mutations | 0 |
| $\mu_D$ | Rate at which $M_A$ alleles will evolve to be expressed | 0 |
| $\beta$ | Coefficient for $F$ overexpression relative to temperature $T$ | 0 |
| $\mu_E$ | Mean expression level for $M_A$ when evolving *de novo* | 0.9 |
| $\sigma_E$ | Standard deviation for $M_A$ expression level when evolving *de novo* | 0.05 |
| $\sigma_F$ | Standard deviation for noise in $F$ expression level | 0.05 |

**Supplementary Table 3:**
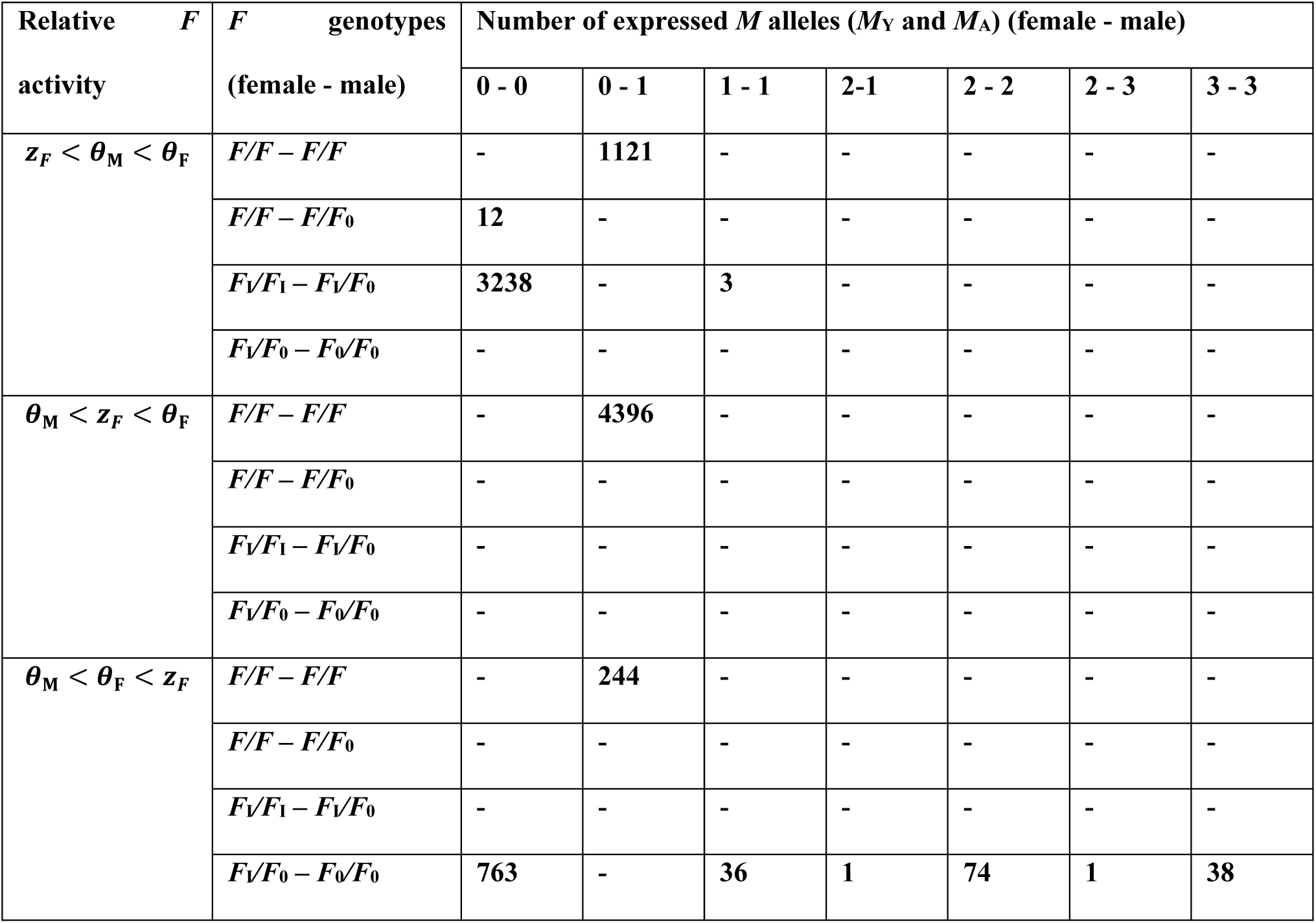
Sex determination systems evolved under different levels of *F* activity relative to the maleness and femaleness thresholds *θ*_M_ and *θ*_F_. Shown are the number of simulations in which a certain combination of female and male genotypes for *F* and *M* were found to be the most prevalent genotype. For *F* genotypes, *F* alleles are indicated by *F* (expressed and sensitive), *F*_I_ (expressed and insensitive), and *F*_0_ (unexpressed). For *M*, we indicate the number of expressed *M* alleles in females and males. Dashes indicate that the system has not been observed under those particular conditions. We performed 10,000 independent simulations under different conditions, of which 9,927 terminated successfully (e.g., did not end with population extinction) and could be categorized.

## Supplementary Figures

**Supplementary Figure 1:**
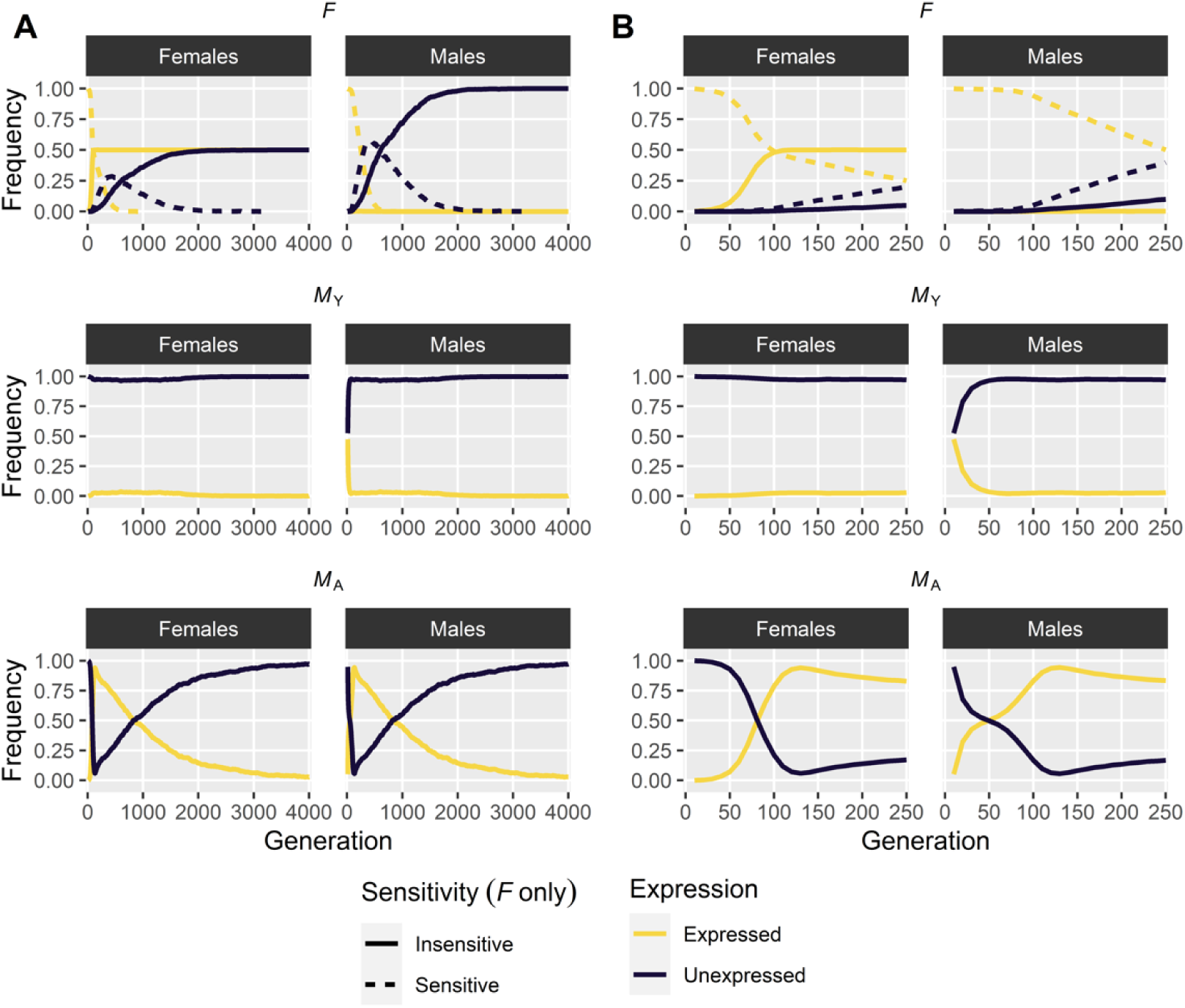
Evolutionary dynamics of sex determination genes. The spread of an insensitive *F* allele enables the fixation of *M*_A_ in both sexes, followed by the gradual accumulation of unexpressed *F* and loss of expressed *M*_A_. (A) Evolutionary dynamics over 5,000 generations. (B) Detail of the initial 250 generations shown in (A). Parameter values: *µ*_D_ = 0.027; *θ*_M_ = 0.399; *θ*_F_ = 0.789.

**Supplementary Figure 2:**
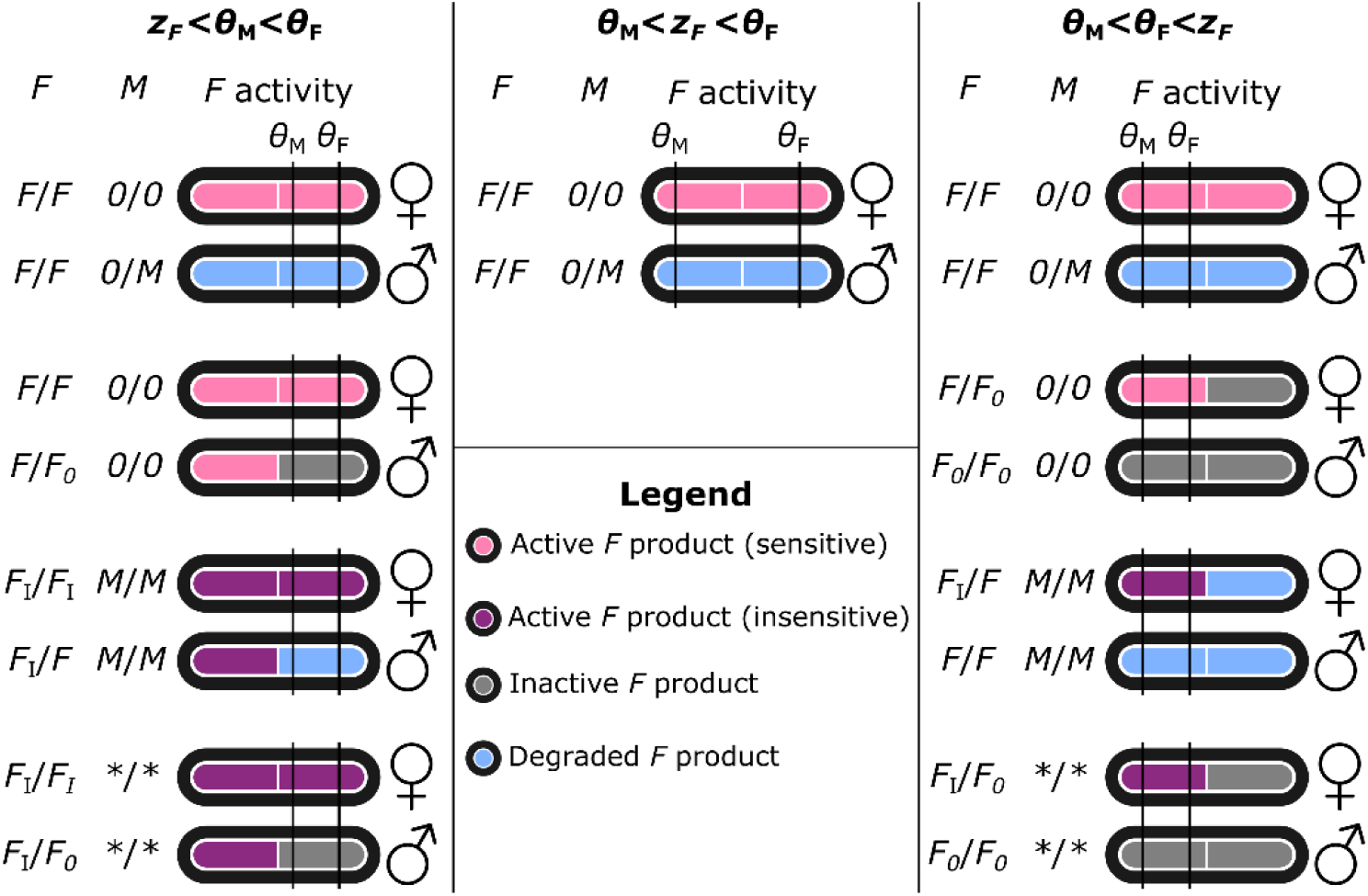
Classification of sex determination systems depending on relative values of thresholds and F activity. Each system is defined by a recurrent pair of female and male genotypes that can only generate those two genotypes (conform ^26^). Here we define three alleles for locus *F*: a regular *F* that is expressed and sensitive to *M*; a variant *F*_I_ that is insensitive to *M* and expressed; and a variant *F*_*0*_ that is unexpressed. For *M*, we distinguish between active (*M*) and inactive (*0*) variants. We use *z*_*F*_ to refer to the activity of a single *F* allele. In systems with both *F*_I_ and *F*_*0*_, *M* has no function and may be present or absent; this is indicated by asterisks (*\*/\**).

**Supplementary Figure 3:**
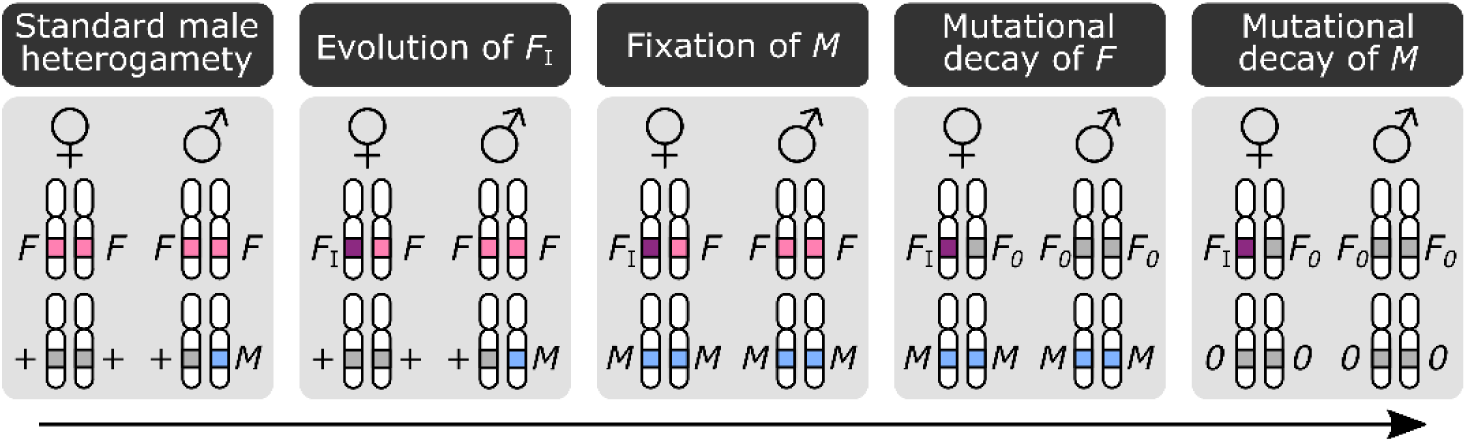
Genetic decay of *F* and *M* following the evolution of an insensitive feminizing F_I_ allele. Starting from a standard male heterogametic system, a feminizing *F*_I_ allele can invade resulting in a transition to a female heterogamety system. For this to occur, an *M* allele must be fixed in both sexes. Fixation of *M* ensures the regularly-sensitive *F* product is always degraded and hence these alleles confer no function leaving them susceptible to genetic decay by mutation accumulation. This can eventually lead to loss of functional *F* in favour of non-expressed *F* alleles (denoted *F*_0_). Loss of *F* obviates the need for *M* to be maintained as no regular F product is generated that must be broken down to ensure maleness in non-*F*_I_-bearing individuals (genetic males). Similar to *F* previously, *M* functionality is no longer necessary and mutations may accumulate by which *M* becomes unexpressed (denoted 0). Note that *F*_I_ is depicted here as a dominant feminizing allele (females *F*_I_//*F*, males *F/F*) but similar scenarios apply for a recessive feminizing *F*_I_ allele (females *F*_I_//*F*_I_, males *F*_I_//*F*). The only difference is that the *F* allele that is susceptible to decay is now only found in males in a heterozygous state rather than in heterozygous females and homozygous males; decay of *F* and *M* can occur according to the same principles as when *F*_I_ is a dominant feminizing allele.

**Supplementary Figure 4:**
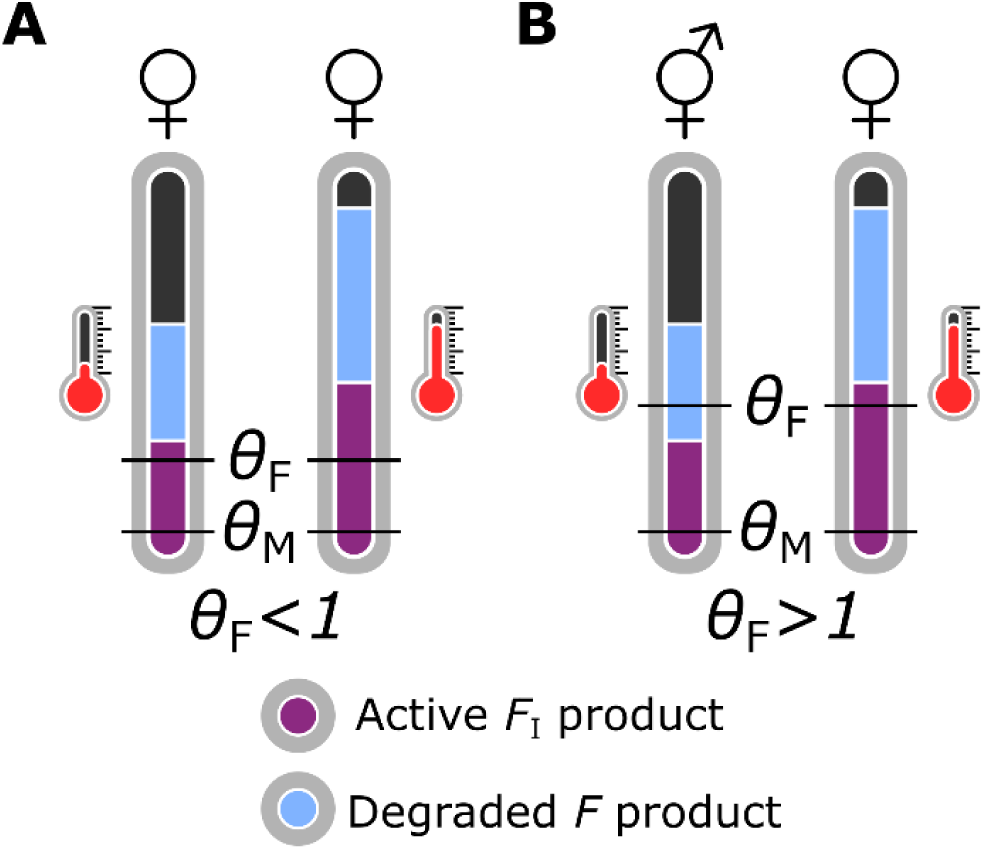
Constraints on the spread of a dominant feminizing *F*_I_ allele. (A) When *θ*_F_ < 1, *F*_I_ can invade in the entire population as a single *F* allele generates sufficient product to induce female development. (B) When *θ*_F_ > 1, *F*_I_ cannot spread in absence of temperature-dependent overexpression as it does not generate sufficient product to induce femaleness, and instead intersexual development is induced in *F*_I_-bearing individuals. At high temperatures, *F* overexpression enables a single *F* allele to be sufficient for feminization, allowing for *F*_I_ to spread. In all cases, we assume a *F*_I_//*F; M/M* genotype so that regular *F* product is degraded.

